# Phasic Neural Stimulation via Frequency-Modulated Kilohertz Signals: An Alternative to Amplitude Modulation

**DOI:** 10.64898/2026.07.15.738601

**Authors:** David Samuel Rose, Aleksandar Opančar, Štěpán Zelníček, Veronika Šromová, Eric Daniel Głowacki

## Abstract

Kilohertz-frequency (kHz) electrical stimulation (1–100 kHz) is emerging as a powerful tool in both invasive and non-invasive neurostimulation applications, including functional electrical stimulation, spinal cord stimulation, and non-invasive brain stimulation. Most commonly these paradigms rely on amplitude modulation (AM-kHz)—achieved via burst or sinusoidal modulation—to produce phasic neural activation. Here, we propose, and validate, an alternative: frequency-modulated kilohertz stimulation (FM-kHz). This approach leverages the distinct strength–frequency dependence of kHz signals, whereby higher carrier frequencies are less efficient in depolarizing neurons than lower frequencies. By sweeping between sub- and suprathreshold frequencies, at a constant amplitude, FM-kHz generates a phasic neural activation envelope analogous to AM-kHz, without requiring amplitude modulation. Using both computational modelling and experimental data from Locusta migratoria (N5 nerve) and the human median nerve, we demonstrate that FM-kHz stimulation: 1. Produces reliable phasic evoked responses at the FM frequency; 2. Enables two degrees of control over stimulation—via FM frequency and frequency deviation. Across models tested, FM-kHz thresholds followed the same increasing strength-frequency relationship as AM-kHz, with FM-kHz requiring modestly higher thresholds at the upper end of the tested frequency range. These findings position FM-kHz as a viable and potentially advantageous alternative to AM-kHz strategies for future neuromodulation devices, and conceptually ground strength-frequency dependence as the key parameter in interpreting the effects of kHz electrical stimulation.

## 1. Introduction

Kilohertz-frequency electrical waveforms are present across a wide range of neuromodulation paradigms, including functional electrical stimulation (FES), transcutaneous peripheral nerve stimulation, interferential current / temporal interference stimulation (TIS), deep brain stimulation, and spinal cord stimulation[1–3]. Although these approaches differ in electrode geometry, invasiveness, and intended neural target, they share a common feature: neural tissue is exposed to rapidly alternating carrier waveforms whose physiological effects depend not only on stimulation amplitude, but also on carrier frequency and low-frequency waveform structure and modulation. In many applications, kHz stimulation is attractive because it may alter recruitment, comfort, selectivity, or perceived sensation compared with conventional low-frequency pulsed-current stimulation [4–8]. Moreover, higher amplitude kHz waveforms can produce a nerve conduction block effect[9,10]. However, despite widespread adoption, the mechanisms by which kHz waveforms excite or modulate neural tissue remain incompletely understood [5,11,12].

Most kHz stimulation paradigms rely on the creation of a low-frequency amplitude envelope via either sinusoidal, ramp, or burst modulation of a kHz carrier [1,13,14]. In interferential current (aka TIS) stimulation, two kHz-frequency currents with a small frequency offset are applied so that their superposition produces an amplitude-modulated waveform at the difference frequency [15,16]. TIS has generated substantial interest as a potential strategy for non-invasive targeting of deeper neural structures[2,17,18]. The intended physiological effect is commonly attributed to the low-frequency envelope rather than to direct activation by the kHz carrier [2,16]. Similar envelope-based logic also appears in premodulated kHz waveforms, in which a single carrier is amplitude-modulated directly, and in burst-based kHz stimulation paradigms used in peripheral and spinal stimulation contexts[19]. These approaches have motivated the idea that neural activation can be driven at the amplitude-modulation frequency, even when the carrier frequency lies in the kilohertz range[20].

This envelope-centred interpretation, however, is incomplete because the kHz carrier itself has frequency-dependent recruitment properties. Their ability to recruit neural tissue depends strongly on carrier frequency, with higher kHz frequencies requiring larger stimulation amplitudes to evoke neural activation than lower kHz frequencies [21,22]. This relationship can be described using the strength–frequency formalism: the threshold current required to excite neural tissue as a function of sinusoidal carrier frequency. Conceptually, this relationship can be viewed as a frequency-domain analogue of the classical strength–duration relationship. Although the strength–frequency relationship has long existed in older biophysical literature on alternating-current excitation [3,23,24], it is largely absent from contemporary discussions of kHz neuromodulation, where emphasis is often placed instead on the low-frequency envelope alone. Reintroducing this formalism is important because it provides a simple way to interpret why different kHz carriers, even with the same nominal amplitude or modulation envelope, may produce different stimulation thresholds and firing patterns.

From this perspective, amplitude-modulated kHz stimulation can be understood as one specific way of driving neural tissue repeatedly above and below the activation threshold. In AM-kHz, the carrier frequency is held fixed while the instantaneous amplitude is periodically increased and decreased (Figure 1a). If the peak amplitude exceeds the activation threshold at that carrier frequency, phasic neural recruitment occurs near the peaks of the amplitude envelope. However, this threshold-crossing logic suggests another possibility. If activation threshold depends on carrier frequency, then a constant-amplitude stimulus could also move above and below threshold by varying the carrier frequency itself. In this case, phasic activation would not arise from an amplitude modulated envelope, but from an excitability envelope that is created as the waveform sweeps between carrier frequencies at a constant amplitude, effectively moving between more- and less-excitable regions of the strength–frequency curve.

**Figure 1.**
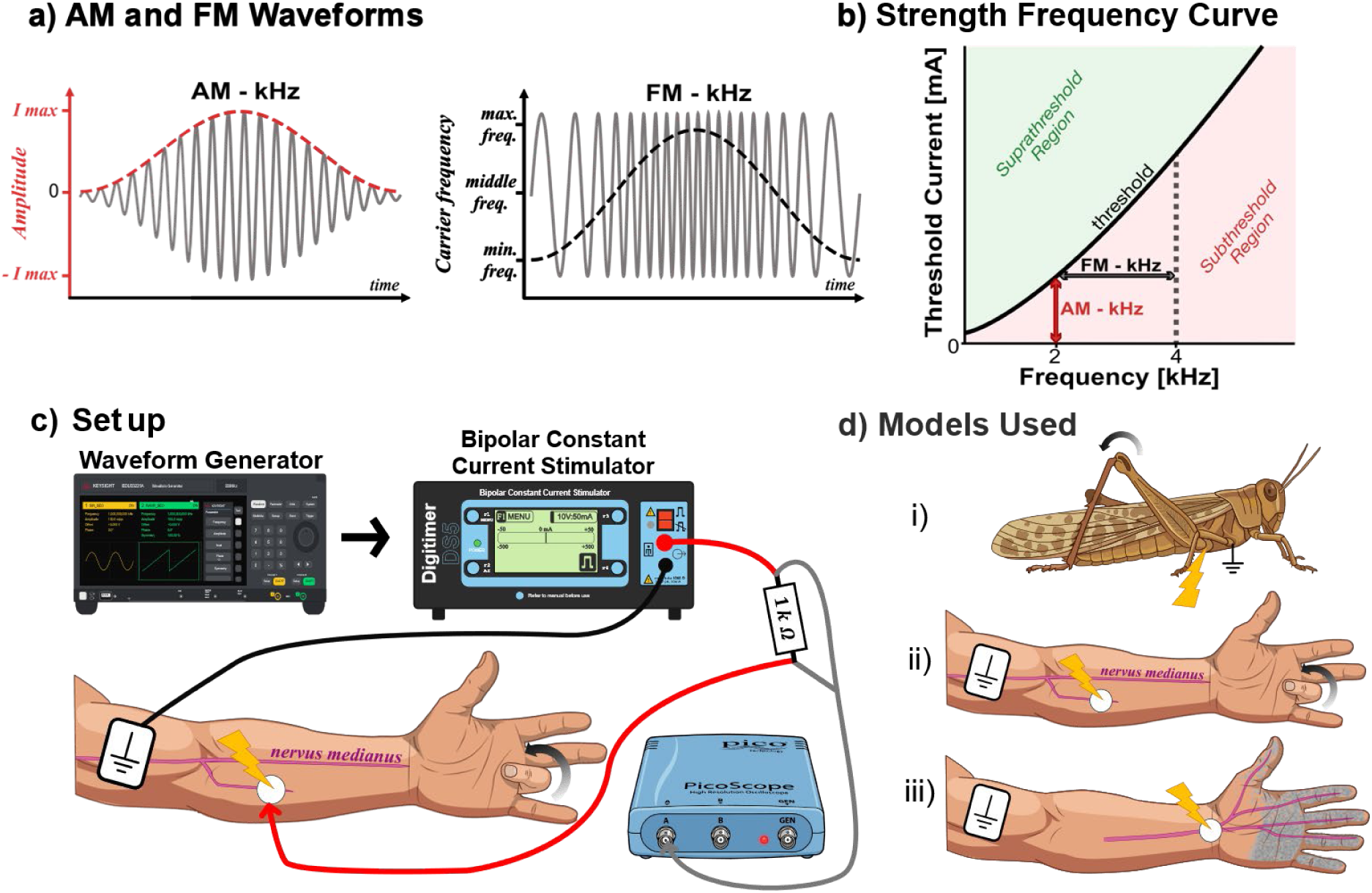
Experimental comparison of AM-kHz and FM-kHz waveforms. a) Schematic of two types of modulation of kHz stimuli; frequency modulation and amplitude modulation. A schematic of each waveform can be seen (grey) along with the modulation signal (red and black). b) Schematic comparison of the two waveforms, AM-kHz vs. FM-kHz, overlaid on a strength–frequency curve. The solid black curve indicates the threshold current required to evoke neural activation at each carrier frequency. Regions below and above this curve indicate subthreshold and suprathreshold stimulus conditions, respectively. In principle, threshold crossing can be achieved either by varying amplitude at a fixed carrier frequency, as in AM-kHz, or by varying carrier frequency at a fixed amplitude, as in FM-kHz. c) Experimental setup of the stimulation circuit used to compare stimulation thresholds between AM-kHz and FM-kHz. Output current from the DS5 bipolar constant current source was measured across a calibrated resistor of known resistance. d) Summary of the three peripheral nerve models used to compare waveforms. i) N5 nerve in the *Locusta migratoria*, ii) Efferent motor nerve in the median nerve innervating the flexor digitorum superficialis, FDS. iii) Afferent median nerve fibres at the carpal tunnel, evoking percepts distally in the palm and fingers.

Here, we test this idea using frequency-modulated kilohertz stimulation (Figure 1a). In FM-kHz, the nominal stimulation amplitude is held constant while the instantaneous carrier frequency is swept over time. The key parameters are modulation frequency (MF), which determines the rate of the frequency sweep, and frequency deviation (FD), which determines the range of carrier frequencies traversed during each modulation cycle. If the lower-frequency portion of the sweep lies closer to threshold and the higher-frequency portion lies farther from threshold, then the same constant-amplitude waveform should preferentially evoke neural activity during the lower-frequency portion of each cycle. In this way, FM-kHz may generate phasic neural activation at the modulation frequency without directly modulating stimulus amplitude (Figure 1b).

This leads to the central question of the present study: can a frequency-modulated excitability envelope produce phasic neural activation comparable to that produced by the amplitude envelope of AM-kHz stimulation? More specifically, can FM-kHz and AM-kHz waveforms be compared under matched conditions, and can their similarities and differences be explained by the underlying strength–frequency relationship? Addressing this question is important not only for evaluating FM-kHz as a potentially useful stimulation waveform, but also for clarifying how kHz stimulation paradigms more generally should be interpreted. Frequency modulation has appeared implicitly in several neuromodulation contexts, including spinal cord stimulation and other kHz-based paradigms, but it has not been defined conceptually in relation to carrier frequency–dependent thresholds. In this study, we compare FM-kHz and AM-kHz stimulation across computational and experimental peripheral nerve models. We first quantify strength–frequency behaviour for both waveforms, then examine how frequency deviation shapes FM-kHz stimulation thresholds. We also assess possible waveform distortions introduced by voltage- and current-controlled stimulation sources, compare evoked sensory percepts during human median nerve afferent stimulation, and use a myelinated axon model to investigate how AM-kHz and FM-kHz differ in their firing dynamics across carrier frequencies and threshold multiples. Together, these experiments test whether FM-kHz can serve as an alternative route to phasic kHz stimulation and whether carrier frequency–dependent thresholds provide a general conceptual framework for interpreting kHz neuromodulation.

## 2. Methods

### 2.1. Ethics

This research was conducted in accordance with the principles embodied in the Declaration of Helsinki and in accordance with local statutory requirements. Experimentation on invertebrates, for example, Locusta migratoria, does not fall under animal experimentation legislation in the Czech Republic (Act No. 246/1992 Sb.). Human experiments were carried out with the written informed consent of each volunteer and were approved by the CEITEC Ethics Committee responsible for research involving healthy human participants. Participants were not monetarily compensated for their time. No adverse events or withdrawals occurred. Human stimulation was delivered with a Digitimer DS5 Isolated Bipolar Constant Current Stimulator, a device that holds clinical approval within the EU.

### 2.2 Electrical stimulation of the locust N5 nerve

Locusts were sourced from a local pet shop and kept in a heat-lamp-equipped terrarium with grass available *ad libitum*. Only adult animals with fully intact anatomy and no visible superficial injuries were enrolled. Anesthesia was induced by cooling each locust to 5 °C for about 15 min. Each animal was then secured ventral-side-up in a custom modelling-clay bed and left to rewarm to room temperature, according to methods outlined previously[25]. For applying stimulation, a constant current bipolar stimulation source, Digitimer DS4, was used to compare AM-kHz and FM-kHz and frequency deviation experiments. For the experiments comparing stimulation sources, both a Digitimer DS4 (current source) and A-301HS amplifier (voltage source; from A.A. Lab systems Ltd.) were used. In both cases, stimulation was delivered via two Ag wires (0.6 mm diameter), with chlorinated tips, placed onto the locust’s cuticle using xyz micromanipulators (S-725CRM, Signatone USA). The Ag wires were chlorinated by immersing them in a standard chlorine-based bleach solution. Electrical contact was improved by placing a drop of ECG gel on each wire tip before it met the cuticle. One wire was placed at the joint joining the hind leg to the thorax as a primary stimulation electrode and a second return electrode was placed centrally on the thorax of the animal. Here, significantly more ECG gel was applied such to decrease the current density such that only the primary stimulation electrode was responsible for determining stimulation threshold. In all instances the output current of both the DS4 and A-301HS were directly measured as the voltage drop across a calibrated 1 kΩ resistor (Picoscope 2206D, input impedance 1 MΩ, software: PicoScope 7 v7.1.39.3737) in series with the stimulation electrode. Every current reported here is one of these directly measured values. The choice of initial waveform, AM-kHz or FM-kHz, was randomized. The ordering of frequency presentation during the collection of the s-f curves for either AM-kHz or FM-kHz was randomized across replicates within a waveform block. Trials were separated by a gap of about 30 s, and the same ramp-to-threshold routine was used for every waveform. Each trial began from the same initial amplitude value. Manual ramping to threshold was used so that the threshold values could precisely be determined. Threshold stimulation, in the locust model, was defined as the minimal phasic stimulation which could be detected. To distinguish sustained phasic activation from onset-only responses, a minimum of five kicks of the leg were required before confirming threshold had been reached. The same threshold definition was used for all locust-based experiments. For each experimental condition measurements were recorded in triplicate.

### 2.3 Electrical stimulation of the human median nerve

Volunteers were enrolled through advertisements placed in the local community and around the institute’s campus. Median nerve motor (efferent) stimulation was performed in 11 participants (mean age 22.6 ± 2.9 years; 27% female), and afferent stimulation in a separate group of 11 participants (mean age 22.3 ± 3 years; 27% female). Participants were not monetarily compensated for their participation.

For both afferent and efferent stimulation in human participants, stimulation was delivered using a Digitimer DS5. Participants were healthy volunteers who provided written informed consent. In all transcutaneous human experiments, commercial gel-assisted Ag/AgCl ECG adhesive electrodes were used as the primary stimulating electrodes. We used much larger return electrodes consisting of gel-assisted carbon TENS electrodes, to purposely reduce current density at the return electrode location, minimizing target stimulation effects. Stimulus waveforms were produced with a two-channel function generator (Keysight Technologies EDU33212A). These voltage signals were supplied to the Digitimer DS5 where the output current was directly measured as the voltage drop across a calibrated 1 kΩ resistor (Picoscope 2206D, input impedance 1 MΩ, software: PicoScope 7 v7.1.39.3737) in series with the stimulation electrode. All currents reported here are these directly measured values. The function generator was driven by a custom Python script that ran the experimental sequence, randomizing trial parameters within each block and keeping the ramping procedure consistent from trial to trial. Participants were kept unaware of the stimulation order, the parameter being varied, and the waveform type, but were not blinded in any other respect. In the comparison of AM-kHz and FM-kHz experiments, the order of which waveform was presented first was randomized. Within each waveform block, frequency presentation order was likewise randomized in software, and successive trials were separated by roughly 30 s. Each experimental condition was measured in triplicate. An identical ramp-to-threshold routine was applied for every waveform across all human sessions.

Beginning at 0.05 Vpp on the Keysight generator, the amplitude was stepped up by 0.05 Vpp every 0.5 s until threshold was detected (or reported by the participant); it was then stepped down by 0.05 Vpp every 0.5 s until the evoked response disappeared, and finally raised again in 0.05 Vpp increments until the response returned. Each trial restarted from this same initial amplitude. The output current ratio for the DS5 was set as 10 V:50 mA. To ensure phasic stimulation was achieved, and not an onset response, often associated with kHz range stimulation, a minimum of five phasic evoked movements/percepts were required to define threshold. This overall protocol was common to every human experiment. Initial waveform selection was randomized, with the experimenter unblinded to which waveform was currently being assessed. However, frequency presentation was also randomized, and the experimenter was blinded to a given frequency presentation, until measuring the threshold current. Participants were blinded throughout the experiment. Within-participant replicates were averaged for a given frequency and waveform before aggregation to group level averages.

#### 2.3.1 Efferent Median nerve stimulation

At the beginning of the experiment, the optimal location for stimulation of the *flexor digitorum superficialis* was found. The final stimulation site was determined optimal when electrode positioning gave rise to the lowest possible current required to elicit a minimal phasic response with a 2kHz amplitude modulated (1 Hz) waveform. To minimize pressure on the forearm—and hence muscle movement—during efferent experiments, participants rested the hand on a cushion with the elbow on the table so that the forearm stayed off the table surface, giving a more stable position. Electrodes were applied to the skin and held in place with kinesiological tape. Threshold was defined as the current required to elicit a consistent phasic response in the evoked finger movement of the participant. Phasic stimulation was defined as the minimal perceivable movement of the participants finger with at least five phasic beats corresponding to the modulation frequency.

#### 2.3.2 Afferent Median nerve stimulation

For afferent nerve stimulation the appropriate stimulation location was determined by placing our gel assisted AgCl electrode along the distal wrist crease, centrally between the two styloid processes at the ends of the ulna and radius. This stimulation site required subjective reporting from the participant, to signal the onset of stimulation, as solely afferent sensory fibres running through the carpal tunnel were stimulated. The evoked sensations were usually present in the palm or fingertips of participants. So, to familiarize participants with the evoked sensations, as well as the onset and disappearance of stimulation, stimulation was verified using a 2kHz amplitude modulated waveform (1 Hz). The participant’s hand was placed flat on the surface of a lightly cushioned table. In all cases, except one, the non-dominant hand of the participant was stimulated. In this one case, due to a minor sports injury in the non-dominant hand of the participant, the dominant hand’s median nerve was stimulated. At each stimulation threshold, while the experimenter was recording the threshold current, the participant was recording where on their palm and fingers they experienced the stimulation. This was achieved via a custom Python script which displayed a standard picture of a palm on a computer monitor [26]. The participant circled, using a mouse, where exactly the evoked sensations occurred. This allowed the quantification of the evoked sensation as well as to determine any trial-by-trial variation that occurred for a set of stimulation parameters.

#### 2.3.3 Quantification of evoked sensory percept maps

During afferent median nerve stimulation, participants were asked to report the onset of stimulation and to indicate the location of the evoked percept. Participants marked the area where they felt the percept on a standardized palmar hand image using a computer mouse [26]. This procedure was repeated for each stimulation condition and replicate, generating a set of participant-drawn percept maps for subsequent spatial analysis.

Participant drawings were exported as individual image files and processed using a custom Python analysis pipeline. Each file name encoded the experimental metadata, including participant ID, waveform type, trial number, carrier frequency, and modulation frequency. The annotation pipeline first rendered each drawing file as an RGB image and then allowed manual annotation of three anatomical landmarks: the wrist centre, middle fingertip, and thumb tip. These landmarks were used to register each participant’s drawing to a common hand template. Handedness was recorded for each file.

Following landmark placement, the perceived stimulation region was traced as one or more closed polygons. Multiple polygons could be drawn for a single trial if the reported percepts were spatially discontinuous. For each annotated file, the pipeline stored metadata, handedness, landmarks, and polygon vertices in a master CSV file.

For quantitative analysis, each participant drawing was spatially aligned to a standardized left- or right-hand template. The three annotated landmarks from the participant drawing were matched to the corresponding three landmarks on the appropriate hand template. A linear spatial registration was computed from these landmark pairs and applied to the participant-drawn percept polygons, placing all reported percept regions into the same template coordinate system for comparison across trials and participants.

After alignment to the hand template, each participant-drawn polygon was rasterized into a binary mask. If a trial contained multiple spatially separate polygons, each polygon was rasterized independently, and the resulting masks were combined using a pixel-wise union. This preserved discontinuous percept regions while ensuring that overlapping pixels were counted only once. The primary percept-size metric was the total number of pixels contained in this combined mask, expressed in aligned template pixels. Left- and right-hand drawings were aligned to their respective templates, and hand side was retained in the output table. For the main area-frequency analysis, hand side was pooled after alignment, while the original handedness remained preserved in the exported data.

Trial-level area values were then grouped by participant, waveform type, hand group, and frequency. Technical replicates were averaged within each participant and stimulation condition, yielding participant-level means. Group-level summaries were then computed across participants for each waveform and frequency. To compare evoked percept area across participants with different absolute percept sizes, each participant’s percept areas were also normalized to that participant’s mean percept area across frequencies within the same waveform condition.

Overlap maps were rendered on the common hand template, with filled regions and outlines used to indicate the spatial distribution of reported percepts.

Final waveform-comparison plots were generated from the participant-level area summary table, group mean of paired participant-level AM–FM comparisons at each frequency. Group plots display mean ± SEM across participants, while paired plots retain individual participant values to show within-participant AM–FM differences.

### 2.4 Computational modelling

Computational modelling in Figs. 6-7 was performed in NEURON using the myelinated mammalian axon model distributed by Mirzakhalili et al.,[5] based on the McIntyre–Richardson–Grill (MRG) framework[27]. The implementation comprises active nodes of Ranvier separated by passive myelinated internodal compartments, including the myelin attachment segment (MYSA), paranode main segment (FLUT), and internode segment (STIN). Nodal membrane dynamics included fast sodium, persistent sodium, slow potassium, leakage, and capacitive currents. Simulations were performed using the 8.7 µm fibre morphology with a total axon length of 100 mm. Channel kinetics and membrane parameters followed the published Mirzakhalili implementation[5], including its modified slow-potassium formulation.

AM-kHz and FM-kHz waveforms were applied extracellularly to the modelled axon in NEURON. Let *t* denote time in seconds, *f_c_* the carrier frequency, *f*_mod_ the modulation frequency (5 Hz), *A* the stimulus amplitude, and *T* = 1/*f*_mod_ the modulation period. Both waveforms were defined so that each modulation cycle began at a low-stimulation boundary. For AM-kHz, a 100% depth nonnegative envelope was used, *e*(*t*) = [1 − cos (2*πf*_mod_*t*)]/2, and the waveform was *I*_AM_(*t*) = *A e*(*t*)sin (2*πf_c_t*). For FM-kHz, the instantaneous frequency was defined as 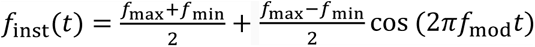, where *f*_max_ = *f*_min_ + 2 kHz, and the waveform was generated as *I*_FM_(*t*) = *A*sin (*ϕ*(*t*)), with 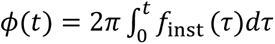. Thus, the AM waveform started at zero amplitude and peaked at mid-cycle, whereas the FM waveform started at its maximal instantaneous frequency and reached its minimum instantaneous frequency at mid-cycle. In the implementation, the FM phase was computed numerically at the simulation time step by discrete integration of the instantaneous frequency.

For direct AM–FM comparisons, *f_c_* in the AM condition was matched to *f*_min_ in the FM condition. Thus, waveforms were compared at the same frequency of peak excitability: AM-kHz reached peak excitability when the envelope amplitude was maximal at *f_c_*, whereas FM-kHz reached peak excitability when the instantaneous frequency was minimal at *f*_min_, with amplitude held constant at *A*.

Simulations were run with a fixed time step in NEURON (*dt* = 0.001 ms; sampling rate 1 MHz). Each batch simulation lasted six modulation cycles, corresponding to 1200 ms at 5 Hz. Activation threshold was defined as the minimum stimulus amplitude that evoked at least one action potential during every modulation cycle for five consecutive cycles. Matched-threshold comparisons were then performed at 1.2×, 1.5×, and 2.0× each waveform’s own phasic activation threshold, so that AM-kHz and FM-kHz firing patterns were compared at equivalent relative stimulus intensities rather than identical absolute amplitudes.

Action potentials were quantified at the root node of the modelled axon using NEURON’s built-in APCount mechanism. In this implementation, the root section corresponds to node 0, with a detection threshold of −20 mV. Representative voltage traces in Figure 6 were displayed from node 0, distal to the stimulation site, to reduce contamination by local kHz-driven membrane oscillations while preserving clearly resolved propagating action potentials. Spikes per cycle were calculated as the mean number of detected spikes per modulation cycle. Burst width was defined within each cycle as the time between the first and last spike assigned to that cycle. Intra-cycle inter-spike interval variability was quantified using only consecutive spike pairs whose two spikes occurred within the same modulation cycle. If *I* denotes the set of such within-cycle intervals, the reported intra-cycle ISI CV was computed as std(*I*)/mean(*I*), using the NumPy population standard deviation and mean.

### 2.5 Statistical Analysis

Statistical analyses were performed using linear mixed-effects models. For threshold experiments comparing AM-kHz and FM-kHz stimulation across carrier frequency, waveform type, and the waveform type × carrier frequency interaction were included as fixed effects. Carrier frequency was treated as a categorical factor because thresholds were measured at discrete frequencies. Locust ID or participant ID was included as a random intercept to account for repeated measurements within the same preparation or participant.

Experimental conditions were measured in triplicate, replicate values were first averaged within each participant or preparation for each waveform and frequency before inferential analysis, unless otherwise stated. Model assumptions were assessed using residual plots, tests for homogeneity of variance, and tests of residual normality. Where raw-measurement models showed heterogeneity of variance or non-normal residuals, the response variable was natural-log transformed before inference. When raw-measurement assumptions were satisfied, analyses were retained on the original measurement scale to aid interpretation.

For the FM-kHz frequency-deviation experiment, frequency deviation was treated as a continuous covariate fixed effect because it represented an ordered numerical variable. Locust ID was included as a random effect.

For evoked percept-area analyses, participant-drawn percept regions were first converted to aligned template pixel areas. Percept area was averaged across replicates for each participant, waveform, and frequency. Absolute percept area was also analysed using a mixed-effects model fitted to ln-transformed mean area, with participant ID as a random intercept and waveform type, frequency, and their interaction as fixed effects. To visualise frequency-dependent changes while reducing between-participant variability in overall percept size, normalised percept area within each participant, and waveform were analysed. The normalised percept-area values were analysed using the same mixed-effects model structure. When a significant main effect of frequency was detected without a significant waveform type × frequency interaction, Tukey-adjusted post-hoc comparisons were performed for frequency levels collapsed across waveform type.

For computational modelling, threshold values and firing metrics were analysed descriptively. Spikes per cycle, burst width, and intra-cycle inter-spike interval coefficient of variation were calculated for each simulated condition and used to compare AM-kHz and FM-kHz firing dynamics across carrier frequencies and threshold multiples. Because these simulations were performed in a single model axon configuration, no population-level inferential statistics were applied to the computational firing metrics.

Unless otherwise stated, statistical significance was assessed at α = 0.05. Group data are reported as mean ± SEM.

## 3. Results

### 3.1 Strength–frequency behaviour for AM- and FM-kHz stimulation

To benchmark the ability of FM-kHz stimulation to achieve phasic suprathreshold activation of peripheral nerve targets compared with AM-kHz stimulation, we conducted current controlled threshold-finding experiments to construct strength–frequency (S–F) curves. These experiments followed the ramping protocol described in Sections 2.2 and 2.3, with AM- and FM-kHz waveforms presented in random order and threshold currents compared across carrier frequencies. In human median nerve afferent stimulation (Figure 2a), human median nerve efferent stimulation (Figure 2b), and locust N5 nerve stimulation (Figure 2c), AM- and FM-kHz waveforms modulated at 1 Hz produced similar S–F curves, with FM-kHz stimulation generally requiring slightly higher threshold currents. The same overall trend was also observed in the MRG neuron model (Figure 2d), where AM- and FM-kHz waveforms showed similar frequency-dependent threshold behaviour.

**Figure 2.**
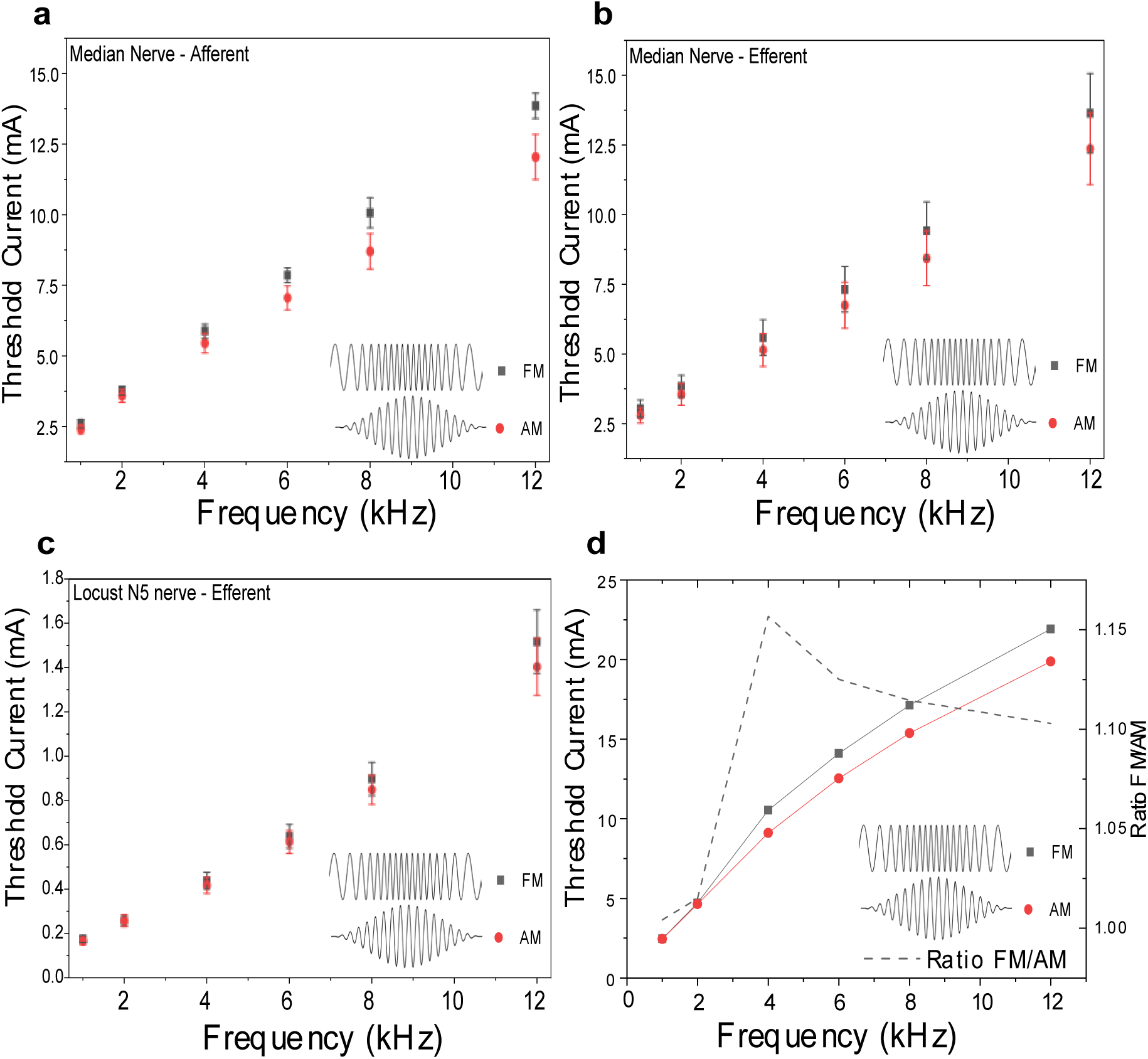
Strength Frequency curves comparing AM-kHz and FM-kHz waveforms. For AM-kHz experiments AMF = 1 and modulation depth = 100%. For FM-kHz MF = 1 Hz and FD = 1kHz. In each model, the threshold current required to elicit the smallest phasic stimulation was employed. For the afferent median nerve model this was titrated to the point where the participant reported phasic onset of a percept in the hand. For the efferent median nerve model and N5 stimulation in the locust, this was titrated to the point of minimum continuous phasic stimulation in the finger/leg. Frequency presentation was randomized for a given waveform, and triplicate technical replicates were collected for each participant. A) Strength frequency curve for afferent median nerve stimulation, at the carpal tunnel (N=11; Error = SEM). B) Strength frequency curve for efferent median nerve stimulation and actuation of the *flexor digitorum superficialis* (N=11; Error = SEM). C) Strength Frequency curve for stimulation of the N5 nerve in the *locusta migratoria* (N=15; Error = SEM). D) Simulated strength frequency curve for the MRG axon model. Here, an AMF = 5 Hz and FD = 1 kHz was used.

For each experimental dataset, threshold currents were analysed using linear mixed-effects models, with waveform, carrier frequency, and their interaction included as fixed effects. Carrier frequency was treated as a categorical factor because thresholds were measured at six discrete experimental frequencies: 1, 2, 4, 6, 8, and 12 kHz. Locust ID or participant ID was included as a random intercept, as appropriate. The appropriate raw data transformations were selected based on model diagnostics: log-transformed threshold current was used when the model showed heterogeneity of variance or non-normal residuals, whereas the raw current values were retained when these assumptions were satisfied.

#### 3.1.1 Locust N5 nerve

For the locust N5 data, raw threshold currents showed increasing variability with carrier frequency. This was supported by significant heterogeneity of variance across waveform-frequency conditions, Levene’s test, p < 0.001, and non-normal residuals from the raw current measurements model, Anderson–Darling AD = 5.703, p < 0.005. Threshold current was therefore natural log transformed prior to inferential testing.

In the log-transformed mixed-effects model, there was a strong main effect of carrier frequency, F(5,154) = 1386.71, p < 0.001, indicating that threshold current increased systematically with increasing carrier frequency. There was also a small but significant main effect of waveform, F(1,154) = 6.64, p = 0.011. Back-transformation of the log-scale waveform effect indicated that FM thresholds were approximately 4.6% higher than AM thresholds overall, corresponding to an estimated FM/AM threshold ratio of 1.046. The waveform × carrier frequency interaction was not significant, F(5,154) = 0.28, p = 0.922, indicating no evidence that the AM–FM threshold difference depended on carrier frequency.

#### 3.1.2 Human median efferent nerve

For human efferent stimulation, the raw average current data satisfied the assumption of homogeneous variance across waveform-frequency conditions, Levene’s test, p = 1.000. Although the corresponding log-transformed data also showed no evidence of heterogeneity, Levene’s test, p = 0.330, the raw current scale was retained as the primary analysis because model assumptions were satisfied and the waveform effect could be interpreted directly in mA.

In the raw-scale mixed-effects model, there was a strong main effect of carrier frequency, F(5,109) = 818.25, p < 0.001, indicating that threshold current increased systematically with carrier frequency. There was also a significant main effect of waveform, F(1,109) = 38.36, p < 0.001. Averaged across the six carrier frequencies and after accounting for the strong effect of carrier frequency, FM stimulation required approximately 0.098 mA higher threshold current than AM stimulation. The waveform × carrier frequency interaction was not significant, F(5,109) = 0.36, p = 0.876, indicating no evidence that the AM–FM threshold difference varied across carrier frequencies. Participant ID accounted for approximately 94.9% of the total model variance, indicating substantial between-participant variation in overall threshold current.

For comparison with the log-transformed locust analysis, the same model was also fitted to natural log transformed threshold current. This gave the same qualitative conclusion: carrier frequency had a strong main effect, F(5,109) = 186.62, p < 0.001; waveform had a significant main effect, F(1,109) = 9.60, p = 0.002; and the waveform × carrier frequency interaction was not significant, F(5,109) = 0.14, p = 0.983. Back-transformation of the log-scale waveform effect indicated that FM thresholds were approximately 7.1% higher than AM thresholds overall, corresponding to an estimated FM/AM threshold ratio of 1.071.

#### 3.1.3 Human median afferent nerve

For human afferent stimulation, the raw average current data showed increasing variability at higher carrier frequencies. This was supported by significant heterogeneity of variance across waveform-frequency conditions, Levene’s test, p < 0.001, and non-normal residuals from the raw-scale model, Anderson–Darling AD = 1.328, p < 0.005. Threshold current was therefore natural log transformed. Following transformation, there was no evidence of heterogeneity of variance, Levene’s test, p = 0.395, and the conditional residuals were consistent with normality, AD = 0.297, p = 0.586.

In the log-transformed mixed-effects model, there was a strong main effect of carrier frequency, F(5,110) = 1121.86, p < 0.001, indicating that threshold current increased systematically with increasing carrier frequency. There was also a significant main effect of waveform, F(1,110) = 57.19, p < 0.001. Back-transformation of the log-scale waveform effect indicated that FM thresholds were approximately 11.8% higher than AM thresholds overall, corresponding to an estimated FM/AM threshold ratio of 1.118. The waveform × carrier frequency interaction was not significant, F(5,110) = 1.27, p = 0.281, indicating no evidence that the proportional AM–FM threshold difference depended on carrier frequency. Participant ID accounted for approximately 80.3% of the total model variance, indicating substantial between-participant variation in overall threshold level.

Together, these analyses show that threshold current increased strongly with carrier frequency across locust N5, human efferent, and human afferent stimulation, for both AM-kHz and FM-kHz. FM-kHz stimulation required slightly higher thresholds than AM-kHz stimulation overall, but there was no evidence in the statistical models that the difference between AM and FM waveforms depended on carrier frequency. Thus, AM- and FM-kHz stimulation produced broadly similar S–F profiles across the models tested.

### 3.2 Frequency-deviation dependence of FM-kHz stimulation threshold

Next, we examined the role of frequency deviation in FM-kHz stimulation using the locust N5 nerve preparation. The modulation frequency was kept constant at 1 Hz, while frequency deviation was varied across six levels from 1 to 10 kHz. The lower bound of the carrier-frequency sweep was fixed at 2 kHz. Threshold current showed a shallow but consistent increase with frequency deviation (Figure 3).

**Figure 3.**
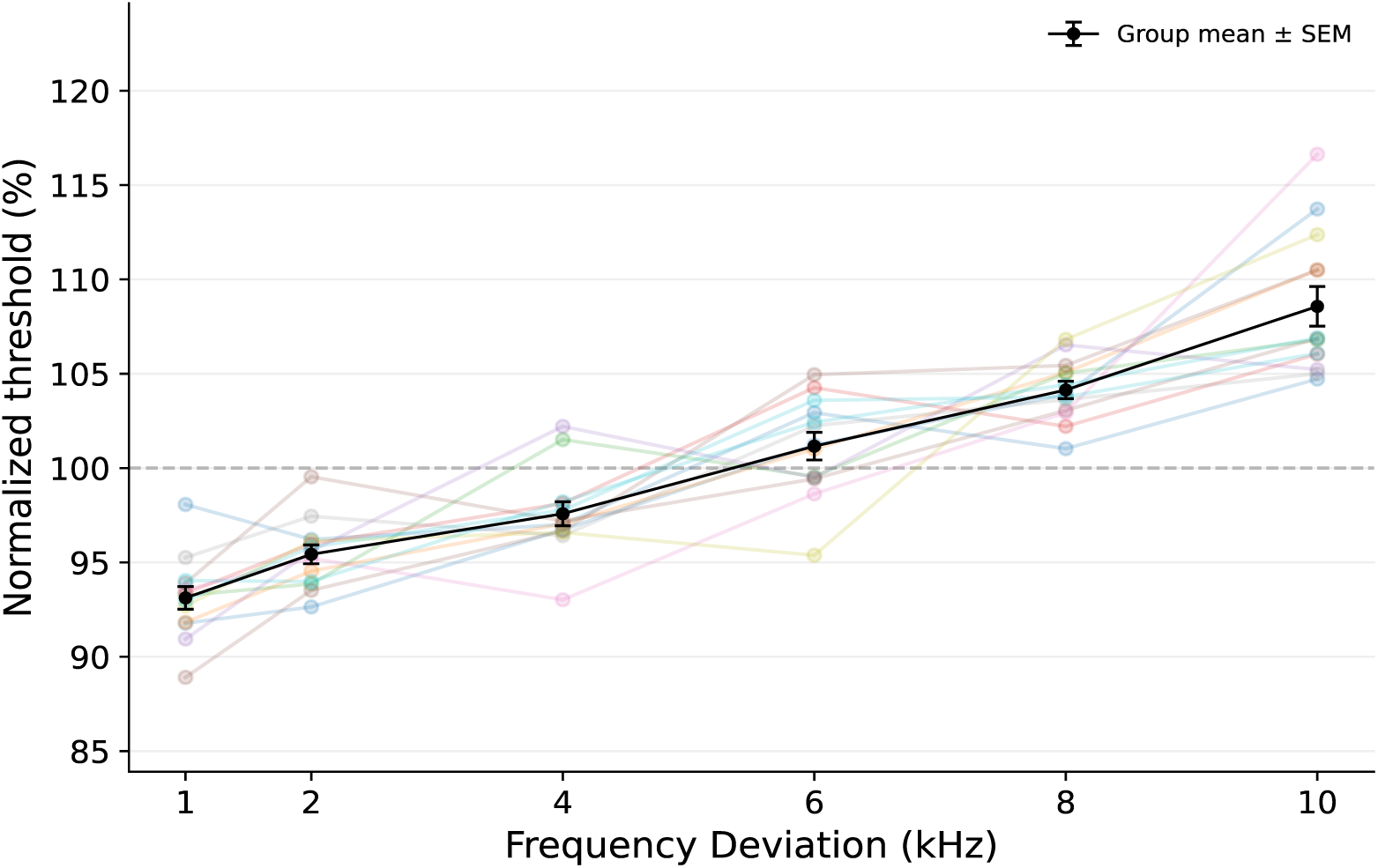
Threshold current required to elicit phasic stimulation as a function of frequency deviation. Here, the threshold required to elicit a phasic stimulation response of the N5 nerve of the *Locusta Migratoria* was used to study the effect of varying frequency deviation (kHz). Increasing the frequency deviation enlarges the range of frequencies through which the waveform is swept during stimulation. MF = 1 Hz and a lower bound of 2 kHz was held constant for all replicates. As such, the total amount of time around the peri-threshold frequencies is reduced with increased frequency deviation resulting in an increased stimulation threshold required to evoke phasic motor stimulation. Individual points show the average normalised threshold values within each locust across replicates, with the group mean plotted across locusts. Thresholds were normalised by dividing each measurement by the average threshold across the tested frequency-deviation range for the corresponding technical replicate. Data are shown as mean ± SEM; N = 13 locusts.

Because frequency deviation was an ordered numerical variable, the primary analysis treated deviation as a fixed covariate. A linear mixed-effects model was fitted to mean threshold current, with frequency deviation included as a fixed effect and locust ID included as a random intercept. The raw mean threshold data showed no evidence of heterogeneity of variance across deviation levels, Levene’s test, p = 1.000, so no transformation was applied. Frequency deviation had a significant positive effect on threshold current, F(1,64) = 243.34, p < 0.001. The fitted slope was 0.00815 mApp per kHz deviation, indicating that threshold current increased as frequency deviation increased. Across the tested range, this corresponded to a fitted increase of approximately 0.073 mApp from the lowest to the highest frequency deviation. Locust ID accounted for approximately 99.0% of the total model variance, indicating large baseline differences in threshold current between preparations.

To express the effect relative to each preparation, the analysis was also repeated using normalised threshold values. The normalised data showed no evidence of heterogeneity of variance across deviation levels, Levene’s test, p = 0.208. Frequency deviation again had a significant positive effect on threshold, F(1,76) = 348.27, p < 0.001. In the normalised analysis, threshold increased by approximately 1.65 percentage points for every 1 kHz increase in frequency deviation. Across the tested range, this corresponded to a fitted increase of approximately 14.8% from the lowest to the highest frequency deviation. Thus, increasing frequency deviation produced a small but consistent increase in the FM-kHz stimulation threshold (Figure 3).

This trend is consistent with the interpretation that larger frequency deviations expose the nerve to a wider range of carrier frequencies during each modulation cycle. If the modulation cycle is kept constant (i.e. 1 Hz) then the total time spent in the peri-threshold region is decreased by including frequencies associated with higher stimulation thresholds. As a result, greater current is required to achieve threshold activation when the frequency deviation is increased.

### 3.3 Voltage- vs. current-controlled stimulation

An important consideration when comparing and implementing FM-kHz stimulation are the stimulation electronics, such as the DS4 and DS5. The load impedance can introduce frequency-dependent waveform distortions[28]. For a voltage-controlled sinusoidal source driving an ideal capacitive load of capacitance *C*, the peak current increases linearly with frequency:

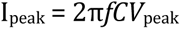

Thus, even if the command voltage amplitude is held constant, the delivered current need not remain constant during a frequency sweep. Although the bulk tissue impedance is often approximated as predominantly resistive in the kHz range, the electrode-tissue interface can retain capacitive and dispersive components. Consequently, under voltage-controlled stimulation, increasing carrier frequency may decrease the effective impedance and thereby increase the current delivered to the sample. This effect is particularly relevant for FM-kHz stimulation, where frequency is deliberately varied over time.

Current-controlled stimulation sources, which are typically the more common choice for neurostimulation contexts, can also introduce a distortion due to the fact that Howland current pump circuits, which comprise these stimulators, suffer from higher frequency amplitude roll-off. Thus, one can expect that when applying FM-kHz waveforms, higher frequencies will have a lower output current. We performed stimulation experiments on locusts with FM-kHz and a 1 Hz modulation frequency and compared current and voltage sources (Figure 4). The different current output FM-kHz “distortion” is clearly visible in both current and voltage-controlled situations (Figure 4a).

**Figure 4.**
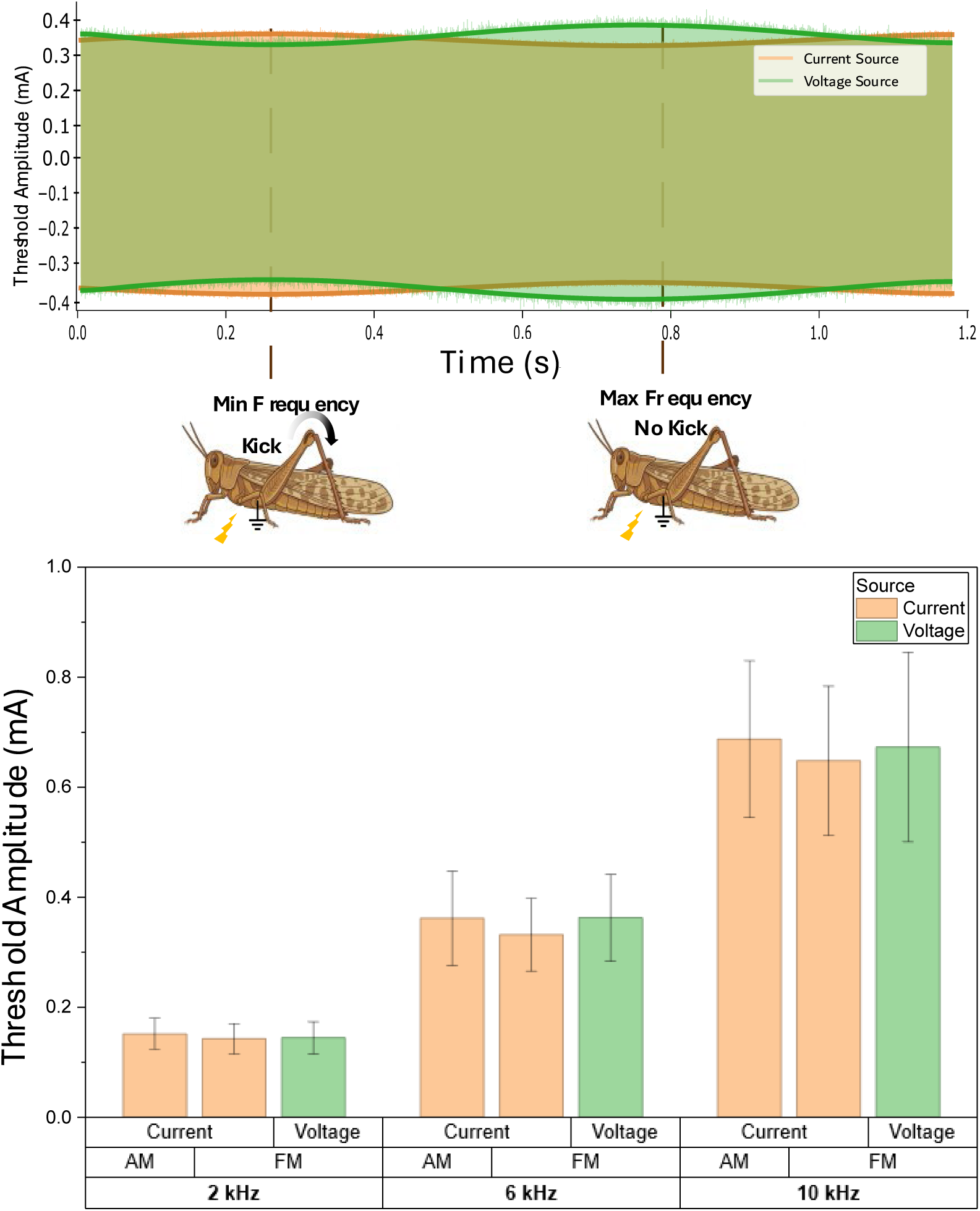
Distortion obtained in waveform amplitude for AM-kHz and FM-kHz. a) Comparison of a constant current source and constant voltage source during FM-kHz. The constant current source amplitude is slightly modulated due to output current roll-off at higher kHz frequencies; this results in a slightly smaller output current at the upper end of the FD range. In contrast, with a constant voltage source, the opposite modulation is observed. Here the lower frequencies have a slightly smaller measured current, largely to do with the frequency dependent impedance properties of the tissue. This results in slightly higher measured currents at the upper end of the FD range. Importantly, the overlayed vertical lines are the observed phase, in the modulation cycle, in which the evoked motor activity was observed. Independent of the slight increases or decreases in output current, the evoked activity was always observed at the lower bound of the FD. b) Here the threshold current for equivalent AM-kHz and FM-kHz waveforms for both a constant current and constant voltage source were determined, for three carrier frequencies. No significant differences were observed in stimulation threshold between voltage and current source, indicating that the slight modulation observed is not the primary contributor to the phasic activation of the FM-kHz. (N=3; Error bars = SEM)

The effect of this on practical stimulation, tested at different centre frequencies, appears to be minor (Figure 4b), as both current and voltage-controlled stimulation experiments yielded threshold values that were insignificantly different. Further, evoked movement occurred during the modulation phase corresponding to the lowest frequency in the FM-kHz sweep in both cases, despite the inverse distortions of the stimulation waveform (Figure 4a). However, when deploying FM-kHz stimulation, these effects should be considered in the context of a given experiment, and true output current should always be measured to properly interpret an experimental result.

### 3.4 Evoked sensory percepts of AM-kHz vs. FM-kHz stimulation

Evoked sensory perception and the topic of percepts evoked by kHz stimulation waveforms is a topic of great interest in the spinal cord stimulation and functional electrical stimulation fields. We set out to quantify if there were differences in the percepts evoked by FM-kHz versus more conventional AM-kHz waveforms when stimulating median nerve afferents at the carpal tunnel. Both produced qualitatively similar evoked percepts in the fingers and palms, with the same phasic activation pattern. We tested as a function of frequency (1-12 kHz) and quantified reported percept region (Figure 5). Both modulation paradigms produced similar percepts with decreasing percept area with increasing carrier frequency.

**Figure 5.**
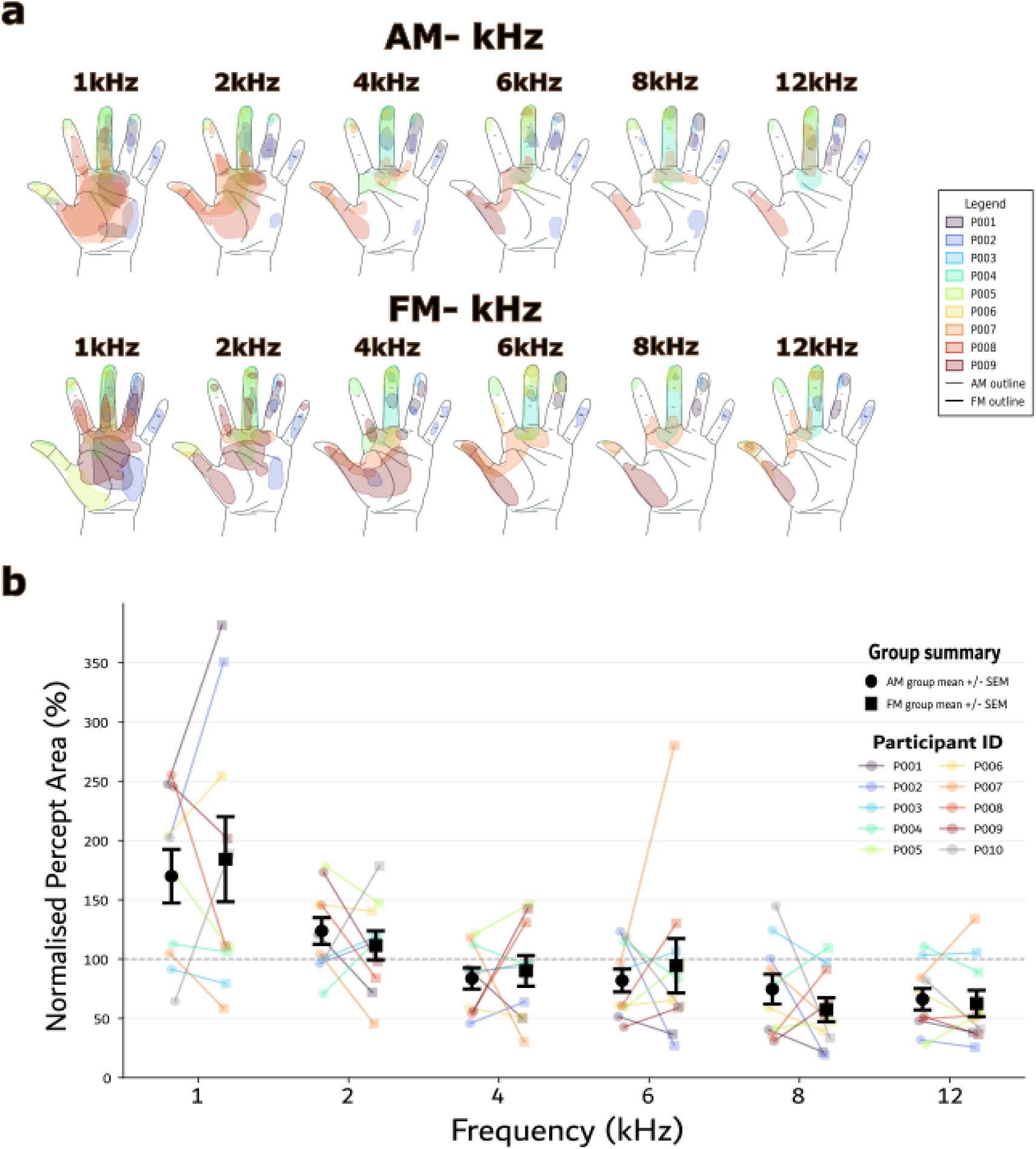
Maps of evoked sensory perceptions induced by afferent median nerve stimulation, for both AM-kHz and FM-kHz, across participants. a) For each replicate, participants marked where they felt the induced sensory percepts. Three technical replicates were collected per frequency and waveform. High inter-replicate consistency was observed for each participant. An apparent decrease in stimulation size appears to follow an increase in frequency in both AM-kHz and FM-kHz. Upper: Hand maps for AM-kHz afferent median nerve stimulation, across participants, AMF = 1 Hz and 100% modulation depth. Lower: Hand maps for FM –kHz afferent median nerve stimulation, FD = 1 kHz and MF = 1 Hz. N=10; Error = SEM. b) Quantification of induced sensory percepts. Normalized percept size was calculated for each participant and waveform by expressing the participant’s mean percept area at each frequency as a percentage of that participant’s average percept area across frequencies within the same waveform. A clear decreasing trend in percept size is apparent as carrier frequency increases. However, no apparent difference between average percept size is obvious between waveforms.

Percept area was first averaged across replicates for each participant, waveform, and frequency. There was relatively low inter-replicate variability for evoked percepts for a given participant and set of waveform parameters. To assess effects on absolute percept area, a mixed-effects model was fitted to ln-transformed mean percept area, with participant ID included as a random factor and waveform type, frequency, and their interaction included as fixed effects. Frequency had a significant effect on ln-transformed percept area, F(5, 99) = 11.54, p < 0.001, indicating that percept area varied across stimulation frequency. The largest percept areas were observed at lower frequencies, with model coefficients indicating higher area at 1 and 2 kHz and lower area at 8 and 12 kHz. In contrast, there was no significant main effect of waveform type, F(1, 99) = 0.00, p = 0.973, and no waveform type × frequency interaction, F(5, 99) = 0.26, p = 0.936. Thus, although percept area changed with frequency, there was no evidence that AM and FM stimulation produced different overall percept areas or different frequency-dependent area profiles. Participant ID accounted for 83.4% of the total variance, indicating substantial between-participant differences in overall percept area.

To visualise frequency-dependent changes while reducing between-participant variability in overall drawing size, percept area was also normalised within each participant and waveform. The data were normalised by dividing by the average percept area across a given replicate’s frequency range, for a participant. These normalised values were then averaged across replicates. A second mixed-effects model was fitted to this normalised percept area measure, again with participant ID as a random factor and waveform type, frequency, and their interaction as fixed effects. This analysis also showed a significant effect of frequency, F(5, 108) = 12.66, p < 0.001. Normalised percept area was highest at low frequencies and decreased at higher frequencies, consistent with the pattern observed in the absolute-area analysis. There was no significant main effect of waveform type, F(1, 108) = 0.00, p = 1.000, and no waveform type × frequency interaction, F(5, 108) = 0.30, p = 0.909. Therefore, the normalised data supported the same conclusion as the absolute-area model: percept area was frequency dependent, but the frequency-dependent pattern did not differ between AM and FM stimulation.

Tukey-adjusted post-hoc comparisons were used to examine the significant main effect of frequency in the normalised percept area model. Normalised percept area was greatest at 1 kHz and was significantly higher than at 2 kHz by 59.4%, p = 0.008; 4 kHz by 90.2%, p < 0.001; 6 kHz by 88.9%, p < 0.001; 8 kHz by 111.0 percentage points, p < 0.001; and 12 kHz by 112.7%, p < 0.001. Normalised percept area at 2 kHz was also significantly higher than at 8 kHz by 51.6%, p = 0.032, and 12 kHz by 53.2%, p = 0.024. No other pairwise frequency comparisons were significant after Tukey correction. Overall, these comparisons indicate that percept area was largest at the lowest frequency and decreased toward higher frequencies.

The underlying mechanism of frequency dependent effects of evoked percept size is not clear from the current data set. Given the high inter-replicate reliability in percept size, it seems unlikely that this is solely explainable due to bias in percept reporting. Instead, it could potentially relate to fibre recruitment patterns as carrier frequency increases. Whether this relates to fewer fibres, or a changing class of fibre, being stimulated requires further investigation.

### 3.5 Computational Modelling

#### 3.5.1 Strength–frequency basis for FM-kHz

To test the mechanistic basis of frequency-modulated kilohertz stimulation (FM-kHz), we first quantified activation threshold as a function of carrier frequency in a single MRG mammalian axon. The central hypothesis was that axonal excitability under kHz sinusoidal stimulation depends strongly on carrier frequency, such that higher frequencies require larger current amplitudes to evoke firing. If so, then phasic recruitment should be achievable not only by modulating amplitude, as in AM-kHz, but also by sweeping frequency across regions of differing excitability along the strength–frequency relationship. For AM-kHz, the stimulation waveform consisted of a kHz sinusoidal carrier with 100% modulation depth at 5 Hz. For FM-kHz, the instantaneous frequency was swept upward by a fixed 2 kHz excursion from the nominal carrier frequency, for example, a 4 kHz carrier corresponded to a 4–6 kHz sweep. In both cases, threshold was defined as the minimum stimulation amplitude that elicited at least one action potential during every modulation cycle for five consecutive cycles. This criterion was chosen to capture reliable phasic entrainment rather than isolated onset firing.

As already shown in Figure 2d, threshold increased with carrier frequency for both AM-kHz and FM-kHz, consistent with the reduced stimulation efficacy of higher-frequency kHz signals. Across the tested range, the two threshold curves remained close, although FM required modestly higher current than AM at intermediate and high carrier frequencies. These results indicate that FM-kHz operates on the same underlying strength–frequency landscape as AM-kHz and can produce reliable phasic activation without modulating amplitude itself. These results closely match the experimental results shown in Figure 2a-c. The threshold curves also explain why FM behaviour should depend strongly on carrier frequency when the FM excursion is held constant. At low carrier frequencies, a 2 kHz sweep spans a relatively large difference in excitability and should therefore approximate AM-like phasic recruitment. At higher carrier frequencies, the same absolute sweep spans a much smaller threshold contrast and should increasingly resemble an effectively unmodulated kHz sinusoid. This analysis therefore establishes both the mechanistic basis for FM-kHz and the normalization required for subsequent comparisons between AM and FM.

#### 3.5.2 Matched-threshold comparison of AM-kHz and FM-kHz firing patterns

Having established that both waveforms operate on the same strength–frequency relationship, we next compared their evoked firing patterns at matched multiples of their own phasic activation thresholds. This normalization was necessary because absolute threshold increased with carrier frequency for both waveforms, and FM thresholds were modestly higher than AM thresholds across most of the tested range. AM and FM were therefore compared at 1.2×, 1.5×, and 2.0× their respective phasic thresholds.

Representative traces are shown in Figure 6. At 2 kHz, AM-kHz and FM-kHz behaved similarly near threshold: both produced one clear burst of action potentials per modulation cycle, with closely matched spike counts and similar temporal structure. At 1.2× threshold, AM generated 169 spikes in total and FM generated 165, with mean burst widths of 51.6 and 47.6 ms, respectively. At 1.5× and 2.0× threshold, AM recruited more spikes than FM, but both waveforms remained clearly phasic and cycle-locked. Thus, at low carrier frequency, a fixed 2 kHz FM sweep traversed a sufficiently large portion of the strength–frequency curve to reproduce the essential phasic behaviour of AM-kHz.

**Figure 6.**
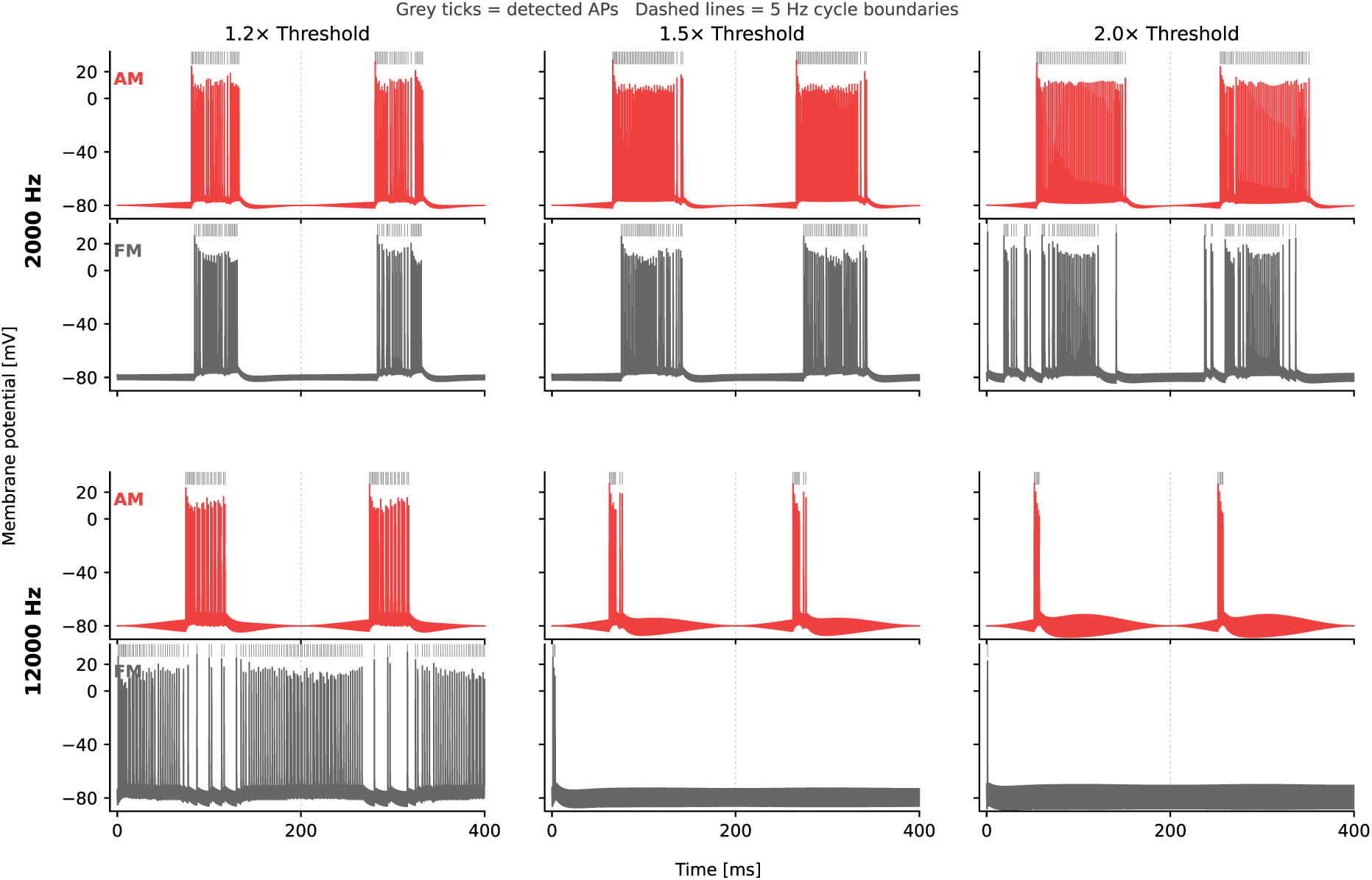
Matched-threshold comparison of AM-kHz and FM-kHz firing patterns in the MRG axon model. Representative membrane-potential traces for AM-kHz (red) and FM-kHz (black) stimulation at carrier frequencies of 2 kHz and 12 kHz, shown at 1.2×, 1.5×, and 2.0× each waveform’s own phasic activation threshold. Grey ticks indicate detected action potentials and dashed vertical lines mark 5 Hz cycle boundaries. FM used a fixed ±1 kHz frequency deviation, with the nominal carrier frequency depicting the lowest bound of this range, whereas AM used 100% modulation depth. Traces are shown from node 0, distal to the stimulation site, to minimize local kHz-driven membrane oscillations while preserving clearly resolved propagating action potentials. At 2 kHz, AM and FM produced broadly similar phasic, cycle-locked bursting. At 12 kHz, AM remained phasic, whereas FM became tonic-like at 1.2× threshold and onset-only at higher threshold multiples.

The comparison changed substantially at 12 kHz. AM remained phasic across all three threshold multiples, although burst duration decreased as amplitude increased. FM, in contrast, entered qualitatively different firing regimes. At 1.2× threshold, the axon no longer produced a compact phasic burst, but instead fired through most of the modulation cycle, yielding a tonic-like response. At 1.5× threshold, the response collapsed to only 3 onset spikes, all in the first cycle, and at 2.0× threshold only a single onset spike remained. This divergence follows directly from the threshold analysis above. A 2–4 kHz sweep spans a relatively steep portion of the strength–frequency curve, allowing FM to move between more- and less-excitable regions during each cycle. By contrast, a 12–14 kHz sweep spans only a small threshold contrast. Once the stimulation amplitude exceeds threshold near the lower end of that sweep, the signal remains near- or above-threshold for much of the cycle, and FM increasingly behaves like an effectively unmodulated high-frequency sinusoid.

#### 3.5.3 Quantitative firing metrics reveal carrier-dependent divergence between AM-kHz and FM-kHz

To quantify the differences observed in the representative traces, we next analyzed three cycle-based metrics: spikes per cycle, burst width, and intra-cycle ISI CV (Figure 7). These measures were chosen to capture complementary aspects of firing behaviour: the strength of recruitment, the fraction of each modulation cycle occupied by spiking, and the temporal regularity of spiking within the active portion of the cycle. The last of these was of particular interest because prior work on sinusoidal and AM-kHz stimulation has shown that inter-spike interval structure can distinguish regular phasic firing from more irregular, quasi-stochastic responses [6].

**Figure 7.**
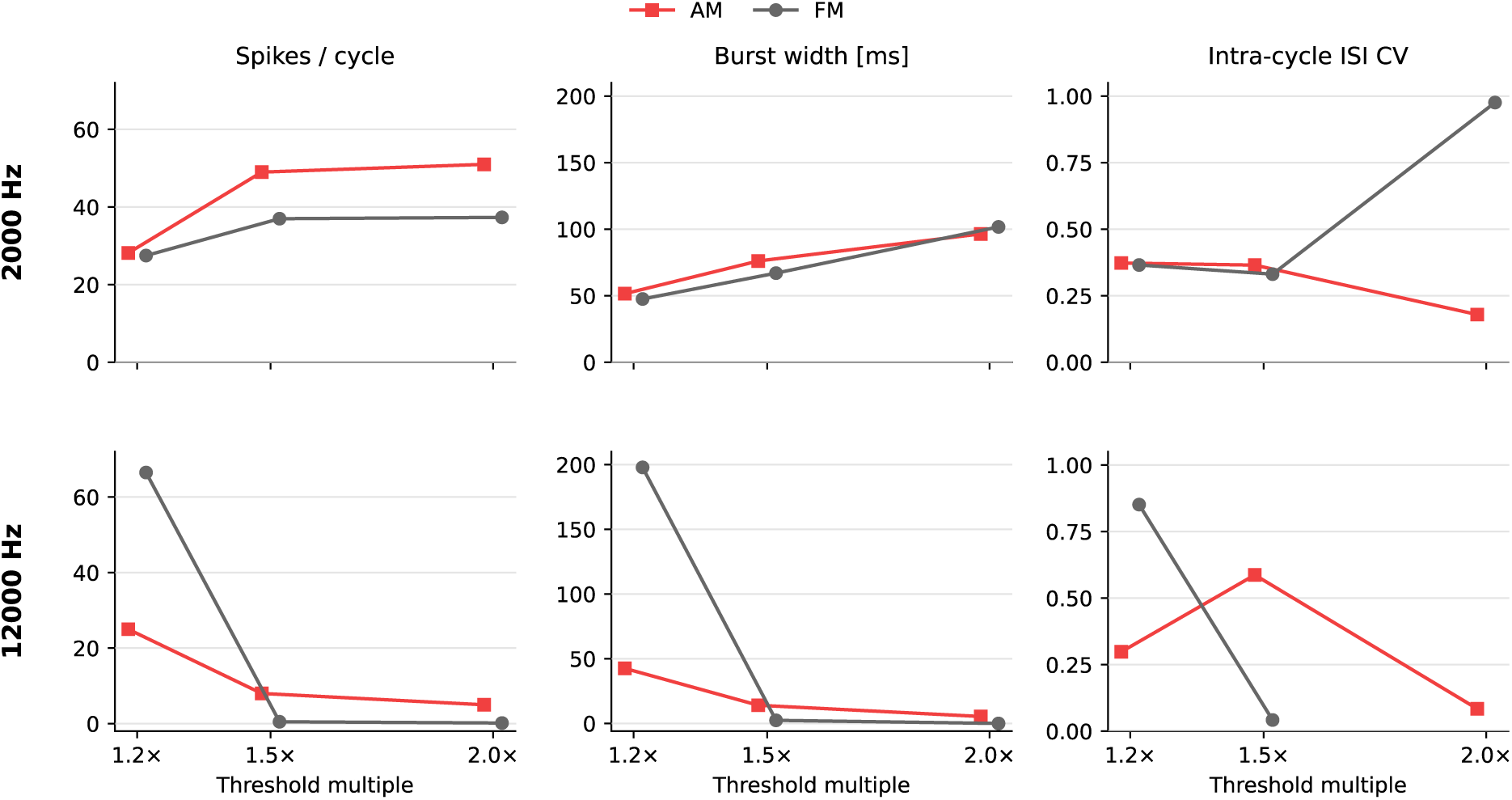
Quantitative firing metrics for AM-kHz and FM-kHz at matched threshold multiples. Summary of the simulations shown in Fig. 3 for AM-kHz (red) and FM-kHz (black) at carrier frequencies of 2 kHz and 12 kHz, evaluated at 1.2×, 1.5×, and 2.0× each waveform’s own phasic activation threshold. Left column: mean number of spikes per 5 Hz modulation cycle. Middle column: mean within-cycle burst width, defined as the time between the first and last spike in each cycle. Right column: coefficient of variation of the intra-cycle inter-spike interval (ISI CV), computed only from spike pairs occurring within the same modulation cycle. At 2 kHz, AM and FM were similar near threshold but diverged at higher amplitudes, particularly in withinburst regularity. At 12 kHz, AM remained phasic across all threshold multiples, whereas FM transitioned from tonic-like to onset-only firing. In the onset-only FM conditions, ISI CV should be interpreted cautiously because it is based on very few spikes.

At 2 kHz, the quantitative metrics confirmed that AM and FM were most similar near threshold. At 1.2× threshold, AM and FM produced 28.2 and 27.5 spikes per cycle, respectively, with burst widths of 51.6 and 47.6 ms and nearly identical intra-cycle ISI CV values of 0.373 and 0.366. Thus, at low carrier frequency and near threshold, the two waveforms were comparable not only in the amount of firing they produced, but also in how that firing was organized within the burst.

At higher threshold multiples, however, the 2 kHz comparison revealed a more subtle divergence. AM produced more spikes per cycle than FM at both 1.5× and 2.0× threshold, while burst widths became relatively similar by 2.0× threshold. The clearest difference emerged in the intra-cycle ISI CV. At 1.5× threshold, AM and FM still showed comparable within-burst irregularity, but by 2.0× threshold AM had become more regular within the burst (ISI CV = 0.179), whereas FM had become markedly more irregular (ISI CV = 0.976). Thus, even in a regime where both waveforms remained clearly phasic at the cycle level, FM no longer matched AM in the fine temporal structure of spiking within each burst. A much stronger separation was observed at 12 kHz. AM remained phasic across all tested threshold multiples, although increasing amplitude progressively compressed the burst in time. FM instead transitioned into entirely different firing regimes. At 1.2× threshold, it produced 66.5 spikes per cycle with a burst width of 198 ms, indicating that spiking occupied nearly the entire 200 ms modulation cycle. The intra-cycle ISI CV in this condition was 0.851, consistent with sustained but irregular tonic-like firing rather than a compact phasic burst. At 1.5× and 2.0× threshold, FM collapsed to onset-only firing. In these onset-dominated conditions, the intra-cycle ISI metric is based on a few spikes and should therefore be interpreted cautiously; here, the loss of a stable ISI structure is itself evidence that sustained phasic activity has broken down.

Taken together, the simulations show that FM-kHz can reproduce AM-like phasic recruitment when the frequency sweep spans a sufficiently large portion of the strength–frequency curve, but that it can also depart into distinct dynamical regimes when the same absolute excursion samples only a shallow portion of that curve. Carrier frequency and frequency deviation are therefore central design parameters for FM-kHz. More broadly, these results provide a mechanistic framework for interpreting the experimental findings presented above in invertebrate and human models.

## 4. Conclusions

Here, we introduce frequency-modulated kilohertz stimulation as an alternative strategy for producing phasic neural activation without directly modulating stimulus amplitude. By exploiting the strength–frequency dependence of kHz stimulation, FM-kHz can drive neural activity through periodic shifts between more- and less-excitable carrier-frequency ranges. Across computational modelling, locust N5 nerve stimulation, and human median nerve afferent and efferent stimulation, FM-kHz produced reliable phasic responses that were broadly comparable to conventional AM-kHz stimulation. The data further show that FM-kHz provides additional waveform-design flexibility through modulation frequency and frequency deviation, while preserving a constant nominal amplitude.

More broadly, these findings provide a conceptual framework for interpreting kHz stimulation in terms of carrier frequency–dependent thresholds. Frequency modulation has appeared implicitly in several neuromodulation paradigms, including spinal cord stimulation, but has lacked a clear formal definition within this context. Here, we show that frequency modulation is best understood through the strength–frequency formalism: an older concept from biophysical stimulation literature that has been largely absent from contemporary discussions of kHz neuromodulation. Under this view, FM-kHz may be useful as a stimulation modality in its own right, but it also serves as a mechanistic lens for understanding how kHz waveforms recruit neural tissue. Specifically, our results indicate that the effects of kHz stimulation should be interpreted not only in terms of amplitude, but also in terms of where the carrier frequency lies on the strength–frequency curve and how much of that curve is traversed during stimulation.

## Code Availabilty Statement

The code used to analyze the stimulation data and run the *PsychoPy* (v2024.2.4) experiments is available on GitHub under the MIT license: https://github.com/dsrose314/AM_FM_CustomCode

## Acknowledgements

This work was supported by the European Research Council (ERC No. 949191), and funding from the Czech Science Foundation GAČR (grant agreement No. 25-18184X), and the BRIDGE project (CEITEC VUT-S-26-9025). The authors acknowledge CzechNanoLab Research Infrastructure supported by MEYS CR (LM2023051).

